# Erythrocyte extravasation drives ferroptosis and inflammatory amplification in eccentric contraction–induced skeletal muscle injury via the heme–HO-1–iron axis

**DOI:** 10.64898/2026.10.21.760722

**Authors:** Menghua Liu, Xiaoyu Wang, Zhengwei Liu, Fengming You, Qi Shen, Miao Zhang, Shumin Chen, Yeteng Jing, Jinsheng Yang, Yingying Wang

## Abstract

Eccentric contraction-induced skeletal muscle injury is a common sports-related condition characterized by pain, impaired muscle function, slow recovery, and frequent recurrence, yet treatment remains symptomatic and mechanism-based targets are lacking. Although microvascular disruption and erythrocyte extravasation characterize severe muscle strain, the metabolic fate of extravasated erythrocytes has received little attention. In hemorrhagic and hemolytic conditions, erythrocyte-derived heme can activate heme oxygenase-1 (HO-1)-mediated iron release and ferroptosis, potentially amplifying inflammation; however, whether this pathway contributes to eccentric muscle injury remains unknown. To address this gap, we established a parameter-controlled rat model in which the gastrocnemius underwent 0, 50, 100, or 150 eccentric stretch repetitions. Muscle torque, gait, weight-bearing, endurance, pain threshold, histopathology, microcirculatory perfusion, macrophage polarization, and heme, ferrous iron, glutathione (GSH), malondialdehyde (MDA), glutathione peroxidase 4 (GPX4), and pro-inflammatory cytokine levels were assessed. Increasing repetitions produced graded functional impairment, myofiber damage, erythrocyte extravasation, inflammatory infiltration, and microcirculatory disturbance. Heme accumulation was accompanied by HO-1 upregulation in perivascular CD68-positive (CD68□) macrophages, muscle iron accumulation, GSH and GPX4 depletion, elevated MDA, and increased M1 macrophage and cytokine levels. These findings suggest a potential heme–HO-1–iron–ferroptosis-associated inflammatory mechanism in eccentric muscle injury, providing a rationale for mechanistic validation and targeted therapeutic development.

## 1. Introduction

Skeletal muscle injury caused by eccentric contraction is one of the most common sports-related injuries [1], clinically characterized by pain, claudication, and reduced muscle strength and endurance [2]. Recovery is typically slow and recurrence frequent, yet current management remains purely symptomatic, with no mechanism-based therapeutic targets available. The pathological basis of this injury has been well characterized: during eccentric contractions, muscle fibers bear high tension while being forcibly lengthened, resulting in Z-disk disruption and myofiber rupture, accompanied by microvascular breakdown, erythrocyte extravasation, and inflammatory edema [3]. However, the downstream metabolic fate of extravasated erythrocytes within the injured tissue-and its pathological consequences-has long been overlooked [4].

In other hemorrhagic and hemolytic conditions, such as intracerebral hemorrhage and hemolytic disorders, a well-defined pathological axis has been established: extravasated erythrocytes undergo degradation with heme accumulation, compensatory upregulation of heme oxygenase-1 (HO-1), and release of free iron, and the resulting iron overload can trigger ferroptosis, as evidenced by elevated free ferrous iron (Fe²□), depleted glutathione (GSH), accumulated malondialdehyde (MDA), and suppressed glutathione peroxidase 4 (GPX4) content [5, 6]. Notably, ferroptosis has rarely been reported in the context of eccentric skeletal muscle injury[7]. Moreover, damage-associated molecular patterns (DAMPs) released during ferroptosis can drive M1 macrophage polarization and the release of pro-inflammatory cytokines such as TNF-α and IL-6, while these cytokines in turn disturb iron homeostasis through autocrine and paracrine mechanisms [8]-suggesting the existence of a self-amplifying “ferroptosis–inflammation” loop that perpetuates tissue damage and impedes repair. Whether a “heme-HO-1-iron overload–ferroptosis” axis is activated after eccentric injury, and whether it couples with M1 polarization to form such an inflammatory amplification loop, remains unknown [9].

Based on these observations, this study hypothesized that eccentric contraction injury activates skeletal muscle ferroptosis via a “microvascular rupture–erythrocyte extravasation–heme/iron overload” axis, and that ferroptotic injury feeds back through M1 macrophage polarization and pro-inflammatory cytokine release to form a self-amplifying inflammatory loop, leading to graded deterioration of motor and sensory function. A parameter-controlled, graded rat model of eccentric contraction-induced injury was accordingly established, and the proposed mechanism was investigated layer by layer, from behavioral assessments and histopathology to in vivo microcirculatory imaging and molecular profiling. The findings suggest activation of a heme–HO-1–ferroptosis-related pathway and its association with M1-driven inflammation, offering a novel mechanistic explanation for skeletal muscle strain and identifying potential targets for mechanism-based intervention.

## 2. Materials and Methods

### 2.1. Animal Housing and Ethical Statement

Male Wistar rats weighing 180 to 220 g were purchased from the Institute of Laboratory Animal Science, Chinese Academy of Medical Sciences. The animals were maintained under a standard 12-hour light–dark cycle with free access to food and water. All animal experiments were reviewed and approved by the Ethics Committee of the Institute of Acupuncture and Moxibustion, China Academy of Chinese Medical Sciences (Beijing, China; approval number: D2024-01-26-05). All procedures were conducted in accordance with the relevant institutional and national guidelines for the care and use of laboratory animals, the ARRIVE guidelines, and the 3Rs principles (replacement, reduction, and refinement). Only male Wistar rats were used in this study. The study was not designed to evaluate sex-specific differences in eccentric contraction-induced muscle injury.

A total of 128 animals were used, with 32 rats per group. The rats were divided into four batches for modeling and intervention. The first batch of 8 rats was used for stimulation parameter optimization, real-time torque recording during the modeling process, and mechanical function retesting after modeling. The second batch of 8 rats underwent behavioral tests without immediate tissue collection, after which these rats were assigned to the following sampling endpoints: 4 rats for Hematoxylin-Eosin (HE) staining and 4 rats for immunofluorescence staining. The third batch of 8 rats was used for speckle (laser speckle) detection and subsequently for lactate measurement (sampled 1 h post-exercise). The fourth batch of 8 rats was used for the remaining biochemical and ELISA assays (sampled 24 h post-exercise).

### 2.2. Establishment of an In Situ Model of Eccentric Contraction-Induced Muscle Strain (ECIMS)

Rats were randomly assigned to four different experimental groups (n = 32 per group). Control group (CON): received anesthesia only, with needle electrodes inserted subcutaneously around the tibial nerve without electrical stimulation intervention. Low-repetition strain group (Model-Low) underwent 50 repetitions of eccentric strain, middle-repetition strain group (Model-Middle) underwent 100 repetitions of eccentric strain, and high-repetition strain group (Model-High) underwent 150 repetitions of eccentric strain. Acute ECIMS was induced on the right gastrocnemius muscle using the Aurora Scientific 1305A in vivo animal experimental system based on optimized mechanical parameters. Briefly, rats were deeply anesthetized with inhaled isoflurane (4% for induction, 2% for maintenance) and placed in a prone position on a temperature-regulated platform. The right knee joint was firmly fixed with a specialized clamp, and the foot was securely attached to a high-speed servo motor force transducer footplate. Needle electrodes were inserted subcutaneously around the tibial nerve to elicit gastrocnemius muscle contractions.

Before formal modeling, the stimulation parameters required optimization: the stimulation current and the stimulation frequency. First, the optimal current was determined at a fixed frequency; then, based on the established optimal current, the optimal frequency was determined to obtain a fully fused tetanic contraction. To determine the optimal stimulation current, tests were conducted using a pulse width of 2 ms with a duration of 60 s. At a fixed frequency of 100 Hz, the current was progressively increased from 0 to 10 mA in step increments of 0.5 mA, with each step maintained for 2.5 s. The optimal current was defined as the lowest current eliciting a maximal, stable tetanic contraction, and was approximately 5.5 mA in the present study. Subsequently, at this optimal current, the optimal frequency was determined by modulating the stimulation frequency from 10 to 130 Hz. The minimum frequency required for complete muscle tetanic contraction was defined as the optimal frequency, which was approximately 100 Hz, with each frequency applied for 30 s.

The acute ECIMS protocol was initiated at the onset of maximal tetanic contraction (optimal current, 100 Hz frequency), with the ankle fixed at 90°. Each single stretch cycle comprised the following: tetanic stimulation was first applied for 200 ms to reach a stable maximal tetanic plateau, after which the servo-motor footplate performed one forced mechanical stretch (from plantarflexion to dorsiflexion) with a stretch angle of 19° and a stretch duration of 50 ms. A strict 30-second rest interval was maintained between consecutive repetitions to allow complete muscle relaxation. The three groups completed 50, 100, and 150 stretch repetitions, respectively.

### 2.3. Gait Analysis

To assess detailed functional changes in gait, three-dimensional gait analysis was performed using the Rodent Gait Analysis System at 24 hours after ECIMS. Three days before the modeling procedure, rats were trained to relieve stress. During each trial, rats traversed a glass corridor illuminated by fluorescent lights, and a camera mounted below the corridor captured dynamic footprints of the paws, from which spatiotemporal gait parameters were reconstructed and quantified automatically. The analyzed parameters included duty cycle, max contact max intensity, max contact mean intensity, print area, and single-stance duration (Stand) of the right (injured) hindlimb. The test was repeated until three consecutive uninterrupted runs were recorded, and the mean values were calculated for each animal. Experiments were conducted in a quiet environment to avoid interference.

### 2.4. Bilateral Weight-Bearing

To monitor non-stimulus-evoked pain behavior changes in rats, the weight-bearing test was performed at 24 hours after ECIMS. A non-invasive dual-plate balance apparatus was used to measure the weight distribution of the hind paws (right side as the muscle-injured side, left side as the control side). Rats were placed in a transparent plastic testing chamber with an inclined plane and forced to stand on their hind paws on independent pressure sensors. Rats were acclimated until they stood quietly with both hind paws symmetrically placed on the two footplates. In normal rats, the force borne by both hind limbs is balanced, but the force on the painful side is significantly reduced. Five separate weight readings from the ipsilateral and contralateral plates were collected from the control panel, with an interval of at least 1 minute between each reading. Results were expressed as the mean ratio of bilateral weight-bearing.

### 2.5. Treadmill Endurance Test

The treadmill endurance test was performed to record the number of electric shocks received by rats when they stopped running due to pain, at 24 hours after ECIMS. Exercise endurance was evaluated on a motorized rat treadmill. Rats were placed in individual lanes with an incline of 15°, and the initial speed was set at 0 m/min with gradual acceleration to 25 m/min over 2 minutes. The cumulative number of electric shocks received within 10 minutes due to the inability to maintain pace from exhaustion or pain was recorded.

### 2.6. Muscle Mechanical Pain Threshold Test (Randall-Selitto test)

The muscle mechanical pain threshold test was performed to detect deep tissue pain threshold at 24 hours after ECIMS. Mechanical nociceptive threshold was assessed using the Randall-Selitto test, which was performed with a digital pressure algometer equipped with a wedge-shaped pusher head. After acclimating the rats to a calm state, a black hood was placed over the rat’s eyes, and mechanical pressure was applied directly to the muscle belly of the right and left gastrocnemius muscles. Measurements were performed at 5-minute intervals to ensure the rats were completely calm, and the average pressure threshold that elicited an immediate paw withdrawal reflex or vocalization was recorded.

### 2.7. HE Staining

At 24 hours after ECIMS, four rats from each group were randomly selected for perfusion fixation, and the right gastrocnemius muscle tissue was harvested. The tissue was fixed in 4% formaldehyde solution, followed by dehydration through a graded ethanol series, clearing in xylene, and paraffin embedding. Transverse sections were cut at a thickness of approximately 5 μm. Sections were routinely deparaffinized and rehydrated, followed by HE staining. Sections were stained sequentially in hematoxylin and eosin for 1 min each, then dehydrated through a graded ethanol series, cleared in xylene, and mounted with neutral resin. Whole-slide digital scanning was performed, and images were collected at 20× magnification. Two sections were selected from each rat for quantitative analysis.

### 2.8. Laser Speckle Contrast Imaging (LSCI)

At 1 hour after ECIMS, rats were anesthetized, and hindlimb microcirculatory blood perfusion was assessed in vivo using a LSCI (Perimed, Sweden). Rats were placed in a prone position with the dorsal aspect of both hindlimbs exposed. Real-time perfusion pseudo-color images were generated, and the mean perfusion unit (PU) and maximum perfusion unit (PU) within the region of interest (ROI) defined over the gastrocnemius muscle belly were quantitatively analyzed. Due to the prone positioning, the left side of the digitally scanned image corresponds to the right hindlimb (injured side).

### 2.9. Immunofluorescence Staining

Four rats from each group were perfused and fixed in 4% formaldehyde for 30 min, then transferred to 0.1 mol/L phosphate buffer (PB) containing 30% sucrose. The following day, frozen sections were prepared. Sections were incubated with a permeabilization agent at 37°C for 2 h, followed by incubation with donkey serum at 37°C for 2 h. Primary antibodies were incubated overnight at 4°C, and secondary antibodies were incubated at 37°C for 2 h. Sections were mounted with DAPI. Primary antibodies used for overnight incubation at 4°C were as follows: anti-CD31 antibody (goat; 1:300; Bio-Techne, USA; AF3628), anti-HO-1 antibody (rabbit; 1:300; Proteintech, China; 10701-1-AP), anti-CD68 antibody (mouse; 1:300; Thermo Fisher, USA; MA5-13324), anti-CD86 antibody (rabbit; 1:300; Proteintech, China; 83213-1-RR). Secondary antibodies used were donkey anti-Goat IgG Highly Cross-Adsorbed Secondary Antibody-Alexa Fluor™ Plus 488 (1:500; Thermo Fisher Scientific, USA; A32814), donkey anti-Mouse IgG Highly Cross-Adsorbed Secondary Antibody, Alexa Fluor™ 488 (1:500; Thermo Fisher Scientific, USA; A-21202), VRDye™ 549 donkey anti-Rabbit IgG Secondary Antibody (1:500; LI-COR Biosciences, USA; 926-54013), DAPI (Ready-to-use) (1:200; Proteintech, China; PR30021), phalloidin-iFluor 405 conjugate (1:200; Thermo Fisher Scientific, USA; A30104CN). For each group, three images from three different rats were selected for statistical analysis.

### 2.10. Biochemical Assays and Enzyme-Linked Immunosorbent Assay (ELISA)

Eight rats from each group were used for biochemical analyses. Lactate levels were determined in serum and gastrocnemius muscle samples collected 1 h after exercise. For all other biochemical analyses, serum and gastrocnemius muscle samples were collected on the second day after exercise (24 h post-exercise). For serum determination, blood samples were collected via cardiac puncture and centrifuged at 3000 rpm at 4°C for 15 min to separate the serum. Immediately after blood collection, the animals were transcardially perfused with pre-cooled physiological saline (0.9% NaCl) via the left ventricle until the outflow from the right atrium ran clear, to remove residual blood from the circulatory system before tissue harvesting. Gastrocnemius muscle tissue was then collected, rinsed with pre-cooled PBS to remove residual blood or other impurities from the surface, weighed, and minced into small pieces. Protease inhibitors (50 μg/mL) were added to the pre-cooled PBS in advance, and the minced tissue was combined with the corresponding volume of PBS (1 g of tissue per 9 mL of PBS) in a glass homogenizer and thoroughly homogenized on ice. The homogenate was transferred to a centrifuge tube and centrifuged at 4°C and 5000 × g for 10 min, after which the supernatant was collected.

Lactate was measured in serum and gastrocnemius muscle samples collected 1 h after exercise using a commercial assay kit (Nanjing Jiancheng, A019-2-2). Fe²□, MDA, GSH, GPX4, heme, TNF-α, and IL-6 were measured in serum and gastrocnemius muscle samples collected 24 h after exercise. Fe²□ was measured using a colorimetric assay kit (Beyotime, S1068S) based on color-complex formation. MDA was measured using a thiobarbituric acid (TBA)-based assay (Nanjing Jiancheng, A003-1). Reduced GSH was measured using a DTNB recycling assay with absorbance measured at 405 nm (Nanjing Jiancheng, A061-5-1). GPX4, TNF-α, and IL-6 were quantified using rat ELISA kits (CUSABIO, EL009869RA, E11987r, and E04640r, respectively), based on antigen–antibody recognition and enzyme-mediated detection. Heme was quantified using a colorimetric assay kit (Abcam, ab272534). All assays were performed according to the manufacturers’ instructions.

### 2.11. Statistical Analysis

All data were analyzed using SPSS 26.0 and visualized using GraphPad Prism 9.0. The Shapiro-Wilk test was used to assess the normality of continuous data distributions, and Levene’s test was utilized to evaluate the homogeneity of variances. Normally distributed data are presented as the mean ± standard error of the mean (S.E.M.), whereas immunofluorescence quantitative data (n = 3) are presented as mean ± standard deviation (SD). For normally distributed variables with equal variances, a one-way analysis of variance (ANOVA) was performed to compare differences among the four experimental groups (CON, M-L, M-M, and M-H), followed by Tukey’s post-hoc test for multiple comparisons. If the assumption of homogeneity of variances was violated, Welch’s ANOVA followed by Dunnett’s T3 post-hoc test was applied. Changes in instantaneous maximal torque across the 0–150 repetition nodes within the M-H group were analyzed using one-way repeated-measures ANOVA. Mauchly’s test was conducted to check the assumption of sphericity, and the Greenhouse-Geisser correction was applied when sphericity was violated. For data that did not conform to a normal distribution, non-parametric tests were employed. A Kruskal-Wallis H test was used to compare overall differences among the four groups, followed by pairwise comparisons of each injury group against the CON group using the Mann-Whitney U test with a Bonferroni correction. For immunofluorescence quantification, due to the small sample size (n = 3) limiting the statistical power of formal normality testing, data were analyzed using one-way ANOVA followed by Tukey’s post-hoc test in accordance with standard histological practices; these results should be interpreted within the context of this small-sample limitation. For all statistical analyses, a two-tailed P < 0.05 was considered statistically significant.

## 3. Results

### 3.1. Establishment and mechanical characterization of a rat eccentric contraction–induced muscle injury model

Using the Aurora Scientific muscle function testing system, electrical stimulation parameters were optimized stepwise to establish a stable and reproducible rat model of eccentric contraction–induced skeletal muscle injury (Figure 1). Incremental current stimulation (0–10 mA; 100 Hz, 2 ms pulse width, 0.5 mA increments every 2.5 s) showed that the evoked torque plateaued at approximately 5.5 mA, a threshold highly consistent across individuals (Figure 1A, B), and gradient frequency stimulation (10–130 Hz) identified 100 Hz as the minimal frequency eliciting maximal tetanic torque (Figure 1C). For injury induction (Figure 1D, E), with the ankle at 90°, stimulation (5.5 mA, 100 Hz) was applied for 200 ms to reach the tetanic plateau, followed by a 50-ms footplate-driven 19° dorsiflexion that imposed an eccentric contraction on the gastrocnemius; this cycle was repeated every 30 s, with rats receiving 0 repetitions (CON), 50 repetitions (M-L), 100 repetitions (M-M), or 150 repetitions (M-H). Maximal tetanic stretch torque declined dynamically during injury, most sharply within the first 100 repetitions (Figure 1F, G), and post-injury reassessment showed that maximal torque at the optimal current, maximal tetanic torque at 100 Hz, and maximal eccentric stretch torque all decreased stepwise with injury severity, whereas the optimal current rose to the 10 mA upper limit and complete tetanus at 100 Hz was lost in all injury groups. The statistical results are presented in Figure 1H, and representative recordings are shown in Figure S1.

**Figure 1.**
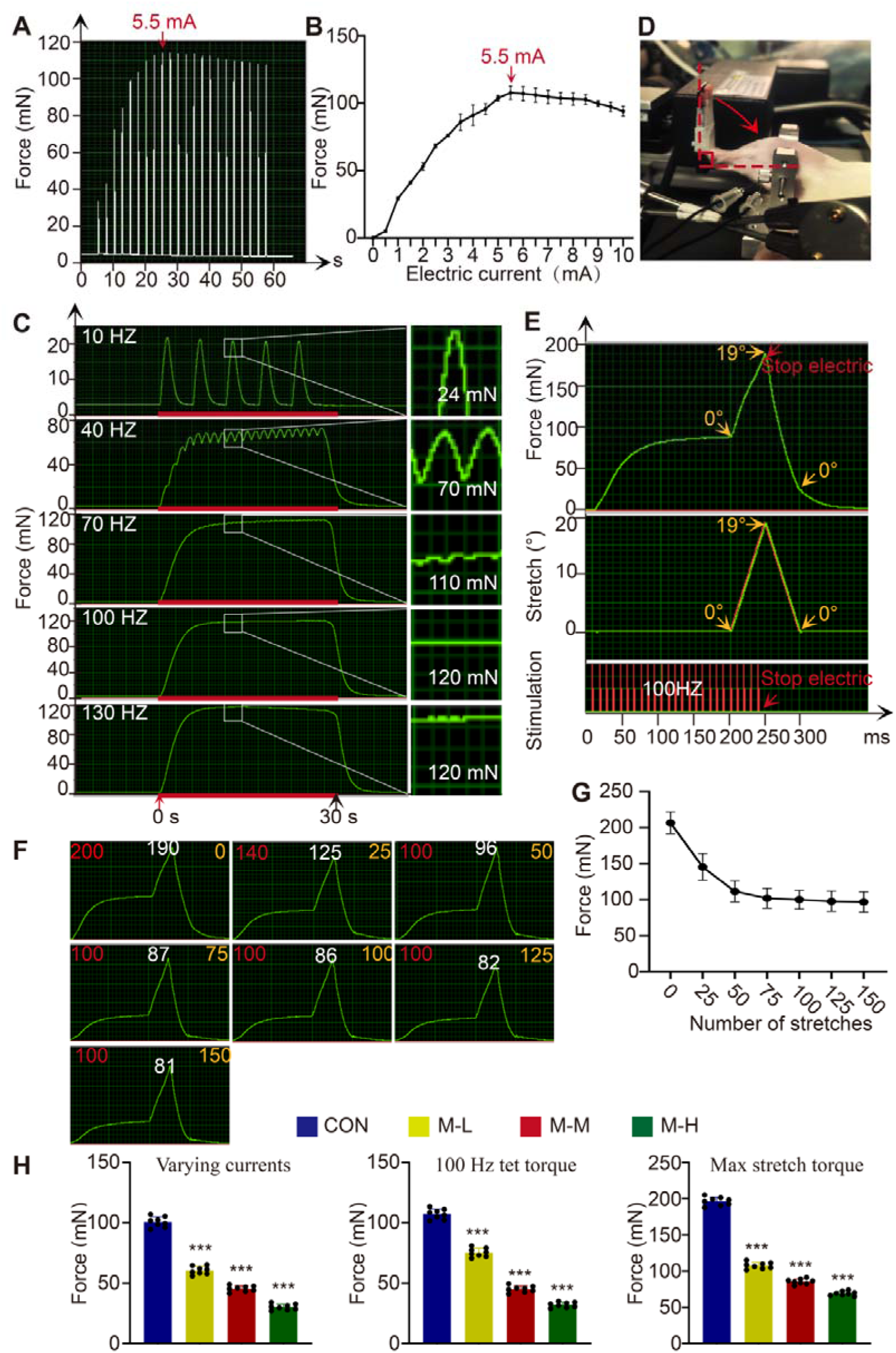
Optimization of stimulation–contraction coupling parameters and dynamic mechanical characterization of a rat eccentric contraction–induced muscle injury model. (A) Time–torque response curve. (B) Current–torque response curve. (C) Tetanic responses of rat muscle at different stimulation frequencies; the red arrow indicates the onset of stimulation and the black arrow its cessation (stimulation duration, 30 s). (D) Photograph of the eccentric contraction–induced strain modeling procedure. (E) Schematic of the injury induction protocol: The figure illustrates a single stimulation cycle. The low, medium, and high-repetition groups (M-L, M-M, and M-H) received 50, 100, and 150 eccentric stretch repetitions. (F) Dynamic decay of maximal torque during eccentric injury in M-H: instantaneous maximal counteracting torque at different repetition nodes (0, 25, 50, 75, 100, 125, 150); red numbers indicate the scale range (mN), white numbers the actual maximal stretch torque (mN), and yellow numbers the stretch repetition count. (G) Summarized curve of maximal muscle torque after different numbers of stretches. (H) Maximal torque at the optimal current, maximal tetanic torque at the optimal current at 100 Hz, and maximal torque during eccentric stretch in each group. n = 8 per group, data are presented as mean ± SEM.

### 3.2. Eccentric contraction injury leads to gait abnormalities, hyperalgesia, and decreased exercise endurance in rats

Having established a mechanically defined, graded injury model, we next evaluated its functional consequences on motor coordination, somatic nociception, and exercise endurance using gait analysis, bilateral static weight-bearing, treadmill endurance, and mechanical pain threshold tests (Figure 2). Three-dimensional gait analysis revealed marked gait disturbances and claudication in injured rats, with progressive deterioration of print area and pressure intensity in the right (injured) hindlimb (Figure 2A); quantitatively, the duty cycle (Figure 2B), max contact max intensity (Figure 2C), max contact mean intensity (Figure 2D), print area (Figure 2E), and single-stance duration (Stand; Figure 2F) of the affected limb all decreased significantly in a repetition-dependent manner, recapitulating the antalgic protective gait observed after clinical muscle injury. Consistently, injured rats shifted weight-bearing from the right hindlimb to the contralateral side (right/left ratio ≈ 1.0 in controls), most severely in the high-repetition group (Figure 2G); received stepwise more electric shocks during the 10-min treadmill test, indicating reduced exercise endurance (Figure 2H); and exhibited a marked, repetition-dependent decrease in the right-hindlimb withdrawal pressure threshold on Randall–Selitto testing, indicating deep-tissue hyperalgesia (Figure 2I). Together, these behavioral data demonstrate that cumulative eccentric injury produces graded impairment of both motor and sensory function.

**Figure 2.**
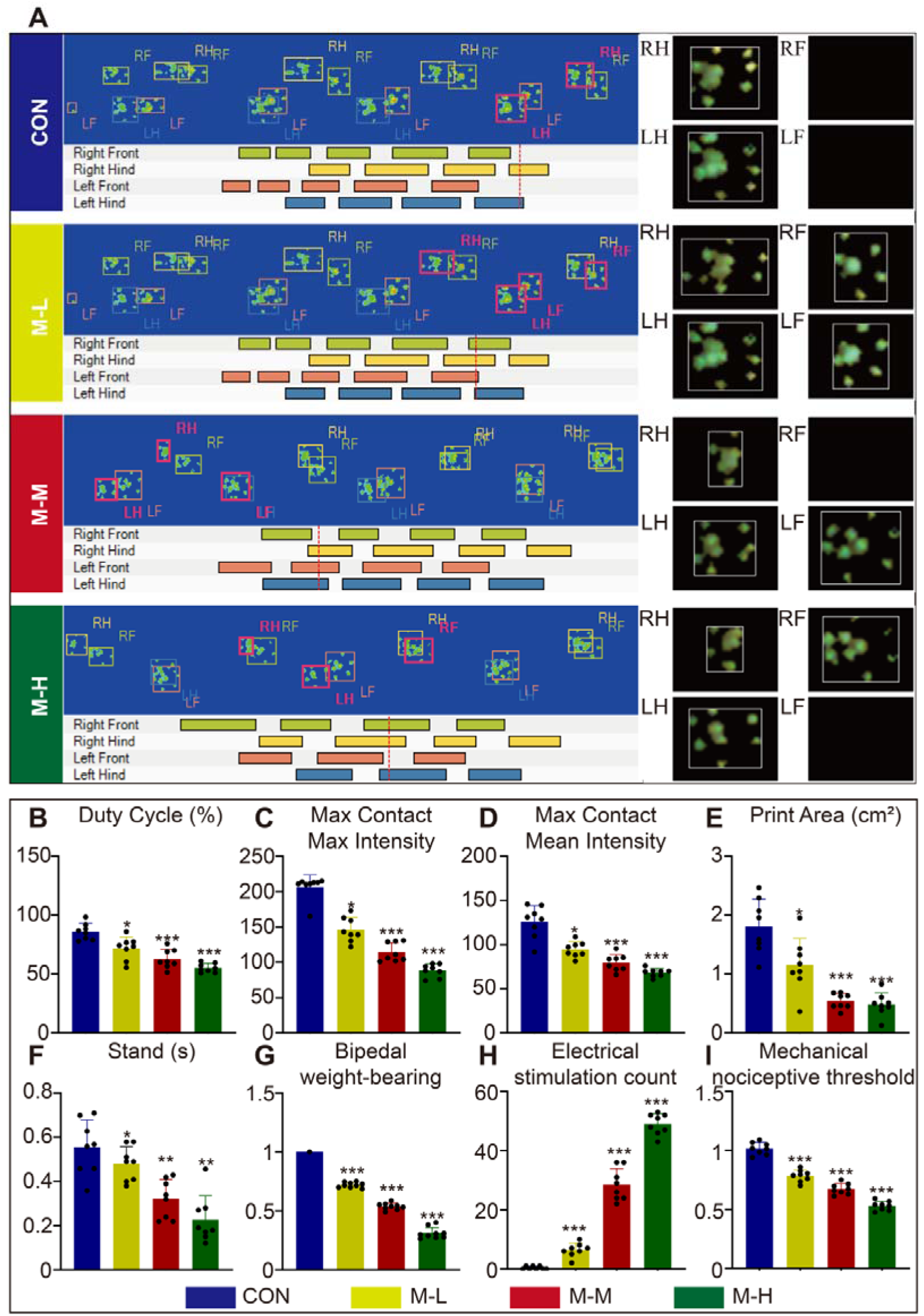
Gait performance, bilateral weight-bearing, exercise endurance, and mechanical pain threshold in rats after eccentric contraction–induced muscle strain. (A) Representative consecutive gait trajectories and pseudocolor topographical maps of plantar pressure distribution in each group. (B-F) Quantification of gait parameters of the right (injured) hindlimb: duty cycle (B), max contact max intensity (C), max contact mean intensity (D), print area (E), and single-stance duration (Stand) (F). (G) Bilateral static weight-bearing distribution between the hindlimbs. (H) Number of electric shocks received during the 10-min treadmill endurance test. (I) Mechanical withdrawal pressure threshold of the hindlimbs measured by Randall–Selitto algometry. n = 8 per group, data are presented as mean ± SEM.

### 3.3. Eccentric contraction injury induces myofiber damage, microvascular hemorrhage, and microcirculatory disturbance

To characterize the histopathological and microcirculatory basis of the functional deficits described above, HE staining and LSCI were performed on the gastrocnemius muscles of all groups (Figure 3). HE-stained sections showed that control muscles contained highly ordered, densely packed myofibers with peripherally located nuclei and no interstitial pathology (Figure 3A), whereas eccentrically stretched muscles exhibited classic, repetition-dependent features of acute injury (Figure 3B): myofiber rupture, distortion, and flocculent necrosis; massive erythrocyte extravasation from ruptured microvessels; and interstitial infiltration of inflammatory cells, with macrophages actively engulfing necrotic myofiber debris. Stereological quantification confirmed that, with increasing repetitions, myofiber cross-sectional area decreased stepwise (Figure 3C) while the interstitial gap widened significantly, peaking in the high-repetition group (Figure 3D), and interstitial erythrocyte and inflammatory cell counts were highest in this group (Figure 3E, F), indicating a substantial local heme burden that provides a morphological basis for the heme–ferroptosis axis examined below. In vivo LSCI (rats imaged prone; the left side of each image corresponds to the injured right hindlimb) showed repetition-dependent expansion and intensification of the high-perfusion signal in the injured region, with mean and maximum perfusion units increasing accordingly (Figure 3G-I), consistent with acute inflammatory hyperemia and microvascular disturbance. Finally, significant lactate accumulation was detected in the gastrocnemius muscle and serum of the M-H group, and marked increases in serum lactate were also detected in the M-L and M-M groups (Figure 3J, K).

**Figure 3.**
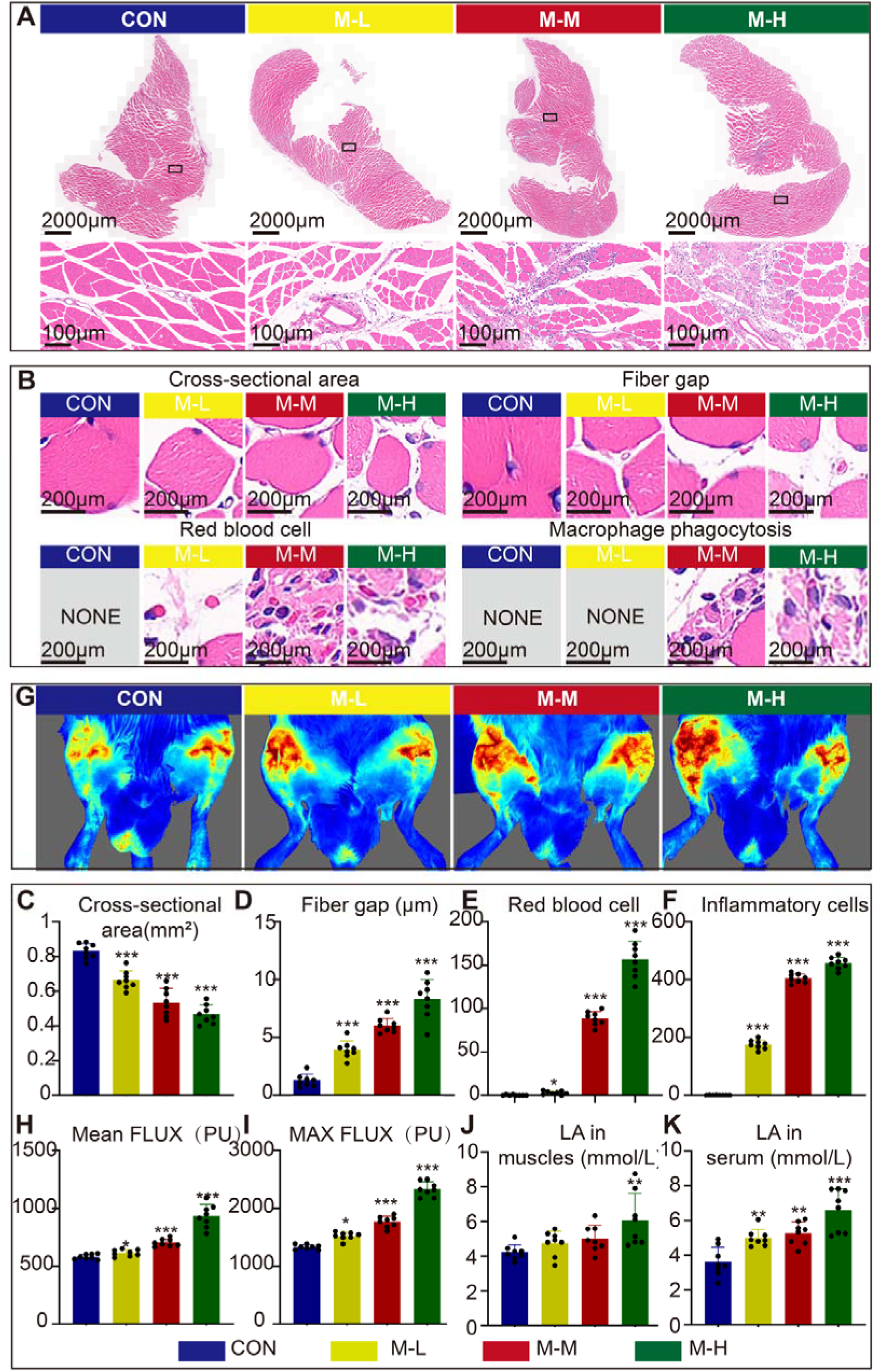
Histopathological damage and in vivo microcirculatory perfusion changes in rat gastrocnemius muscle after eccentric contraction–induced strain. (A) Panoramic scans (scale bar, 2000 μm) and 20× magnified images (scale bar, 100 μm) of HE-stained gastrocnemius sections from each group. (B) Representative histopathological features of eccentric injury (scale bar, 200 μm): reduced myofiber cross-sectional area, widened interstitial fiber gap due to edema, erythrocyte extravasation from ruptured microvessels, and macrophage phagocytosis of necrotic myofibers; “NONE” indicates that the corresponding feature was absent in that group. (C–F) Quantification of myofiber cross-sectional area (C), interstitial fiber gap (D), extravasated erythrocyte count (E), and infiltrating inflammatory cell count (F) in each group. (G) Representative in vivo LSCI pseudocolor perfusion maps of both hindlimbs (rats imaged in the prone position; the left side of each image corresponds to the injured right hindlimb). (H, I) Quantification of mean (H) and maximum (I) blood perfusion (in perfusion units, PU) in the right hindlimb muscle region. (J, K) Lactate (LA) content in the gastrocnemius muscle (J) and serum (K) of each group. n = 8 per group, data are presented as mean ± SEM.

### 3.4. Cumulative injury induces heme accumulation and perivascular macrophage HO-1 upregulation

Given the substantial erythrocyte extravasation and microvascular disruption observed histologically (Figure 3), we next investigated the downstream fate of extravasated erythrocytes by measuring heme, a major product of erythrocyte breakdown, in the serum and gastrocnemius muscle (Figure 4). Heme content was significantly elevated in both the serum and gastrocnemius muscle of the injury groups compared with controls, with higher levels observed in muscle tissue than in serum (Figure 4A, B). We then examined heme catabolism by multiplex immunofluorescence staining for heme oxygenase-1 (HO-1), the rate-limiting heme-degrading enzyme, together with CD31 and phalloidin (Figure 4C). HO-1-positive signals were concentrated in perivascular regions (white boxes) and were markedly weaker in vessel-distant muscle regions (red boxes). The number of HO-1-positive cells was markedly higher in perivascular regions than in distant parenchyma (Figure 4D, E), and the red-box-to-white-box ratio further confirmed this predominant perivascular localization (Figure 4F). HO-1/CD68/DAPI co-staining further revealed clear co-localization of HO-1 with the macrophage marker CD68, with the number of HO-1□/CD68□ double-positive cells increasing significantly with injury severity (Figure 4G, H). These results indicate that cumulative injury promotes erythrocyte extravasation and heme accumulation in both the circulation and injured muscle, with greater heme accumulation in muscle tissue, accompanied by HO-1 upregulation predominantly in perivascular macrophages.

**Figure 4.**
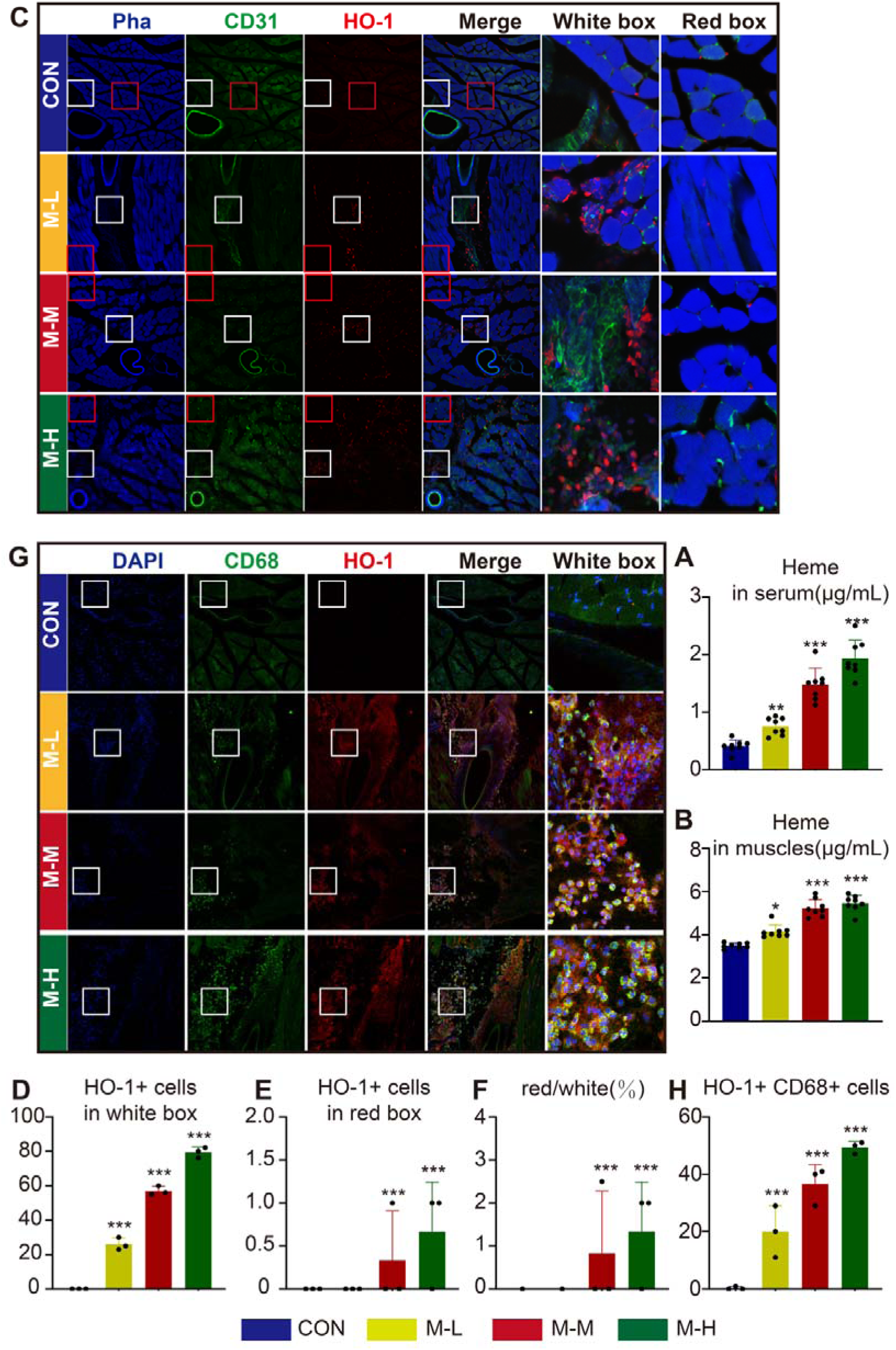
Heme accumulation and perivascular HO-1 upregulation in macrophages after eccentric contraction–induced muscle strain. (A, B) Heme content in serum (A) and gastrocnemius muscle (B) of each group. (C) Representative multiplex immunofluorescence images of phalloidin (Pha, blue), CD31 (green), and HO-1 (red) in the right gastrocnemius muscle of each group; white boxes indicate perivascular regions and red boxes indicate vessel-distant muscle regions, with magnified views shown in the rightmost two columns. (D, E) Quantification of HO-1-positive cells in perivascular white-box regions (D) and in vessel-distant red-box regions (E). (F) Ratio of HO-1-positive cells in red-box versus white-box regions (red/white, %). (G) Representative immunofluorescence co-localization images of DAPI (blue), CD68 (green), and HO-1 (red) in the gastrocnemius muscle of each group, with magnified views of white-box regions in the rightmost column. (H) Quantification of HO-1□/CD68□ double-positive cells in each group. n = 8 per group (A, B), data are presented as mean ± SEM. n = 3 per group (D-H), data are presented as mean ± SD.

### 3.5. Erythrocyte extravasation is associated with muscle iron accumulation and a ferroptosis-related molecular phenotype

The perivascular upregulation of HO-1 (Figure 4) suggests enhanced heme degradation and consequent local iron release. Meanwhile, the staining in Figure 3B revealed necrotic changes in muscle fibers surrounding macrophages. The iron released through HO-1-mediated heme degradation may contribute to local iron accumulation and ferroptosis-related injury in the surrounding muscle tissue. Consistent with this hypothesis, free Fe²□ levels were significantly elevated in the gastrocnemius muscle in the M-M and M-H groups and in serum in all injury groups (Figure 5A, B). Core ferroptosis-related markers showed a coordinated pattern predominantly in skeletal muscle. GSH levels were significantly decreased in muscle, with a more pronounced reduction observed with increasing repetition number (Figure 5C), whereas no statistically significant differences in serum GSH levels were observed among the groups (Figure 5D). Similarly, MDA was significantly increased in muscle across all injury groups and in serum in the M-H group (Figure 5E, F). In contrast, GPX4 content was significantly decreased in M-M and M-H muscle (Figure 5G), whereas no statistically significant difference was observed in serum GPX4 content (Figure 5H). Collectively, these findings indicate that eccentric contraction-induced muscle injury is associated with a ferroptosis-related phenotype predominantly in skeletal muscle, characterized by local iron accumulation, muscle GSH depletion, enhanced lipid peroxidation, and reduced muscle GPX4 content.

**Figure 5.**
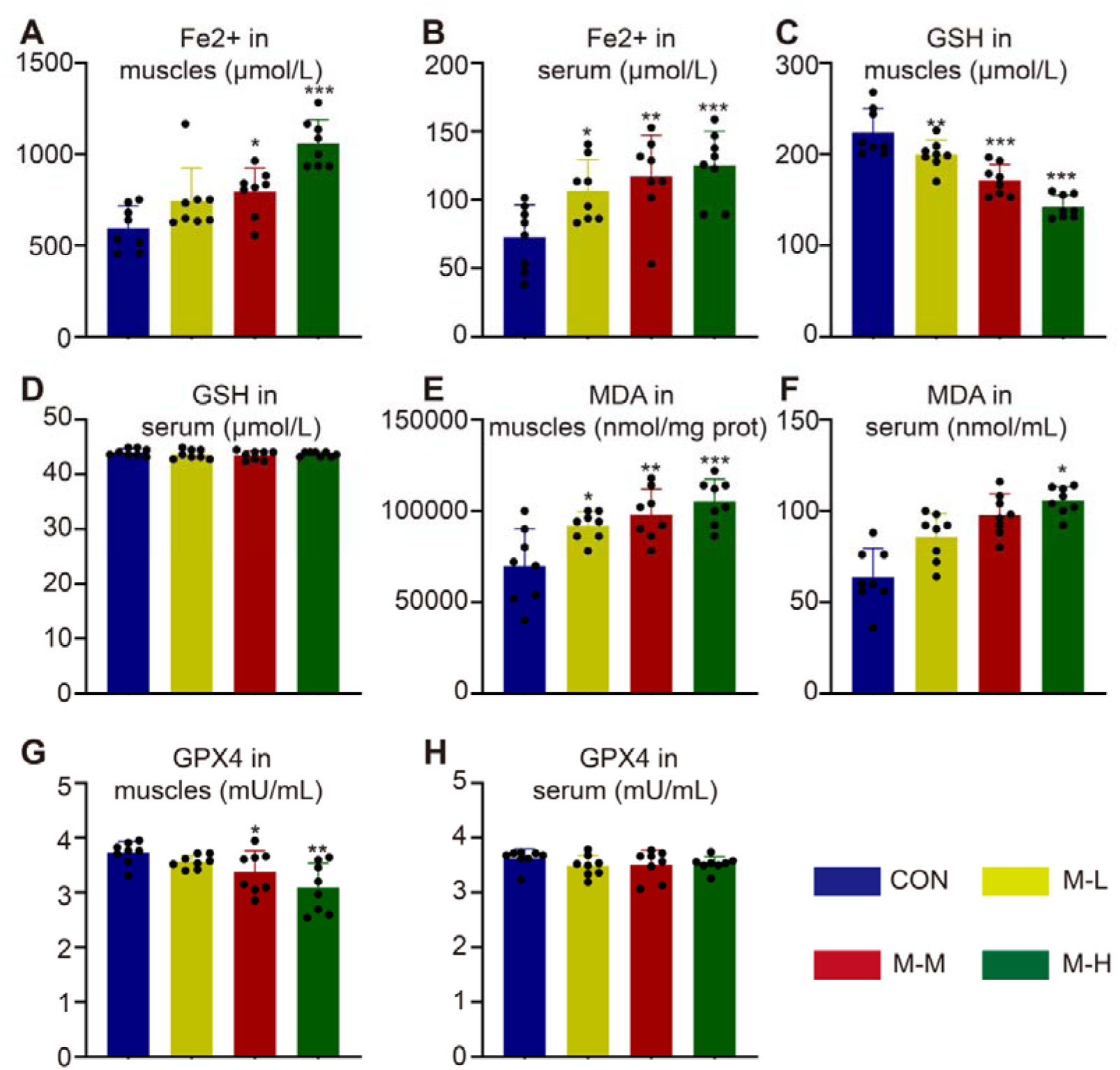
Changes in core ferroptosis markers in rats after eccentric contraction–induced muscle strain. (A, B) Free Fe²□ content in the gastrocnemius muscle (A) and serum (B) of each group. (C, D) Glutathione (GSH) content in the gastrocnemius muscle (C) and serum (D). (E, F) MDA content in the gastrocnemius muscle (E) and serum (F). (G, H) GPX4 content in the gastrocnemius muscle (G) and serum (H). n = 8 per group, data are presented as mean ± SEM.

### 3.6. Eccentric injury is accompanied by M1 macrophage polarization and pro-inflammatory cytokine elevation

Because iron overload and lipid peroxidation can propagate inflammation, we finally examined how the ferroptotic signature identified above (Figure 5) relates to the post-injury inflammatory cascade; to this end, macrophage polarization in the gastrocnemius muscle was assessed by CD68/CD86/phalloidin immunofluorescence co-staining (Figure 6A). Infiltration of CD68□CD86□ double-positive M1 pro-inflammatory macrophages increased significantly and repetition-dependently in the injury groups (Figure 6B), indicating a pro-inflammatory phenotypic shift in the local microenvironment. Consistently, TNF-α was significantly elevated in both the gastrocnemius muscle and serum in all injury groups (Figure 6C, D). IL-6 was significantly elevated in muscle in the M-M and M-H groups and in serum in all injury groups (Figure 6E, F). These findings are consistent with a potential ferroptosis-associated inflammatory feedback loop, although causal mediation remains to be established.

**Figure 6.**
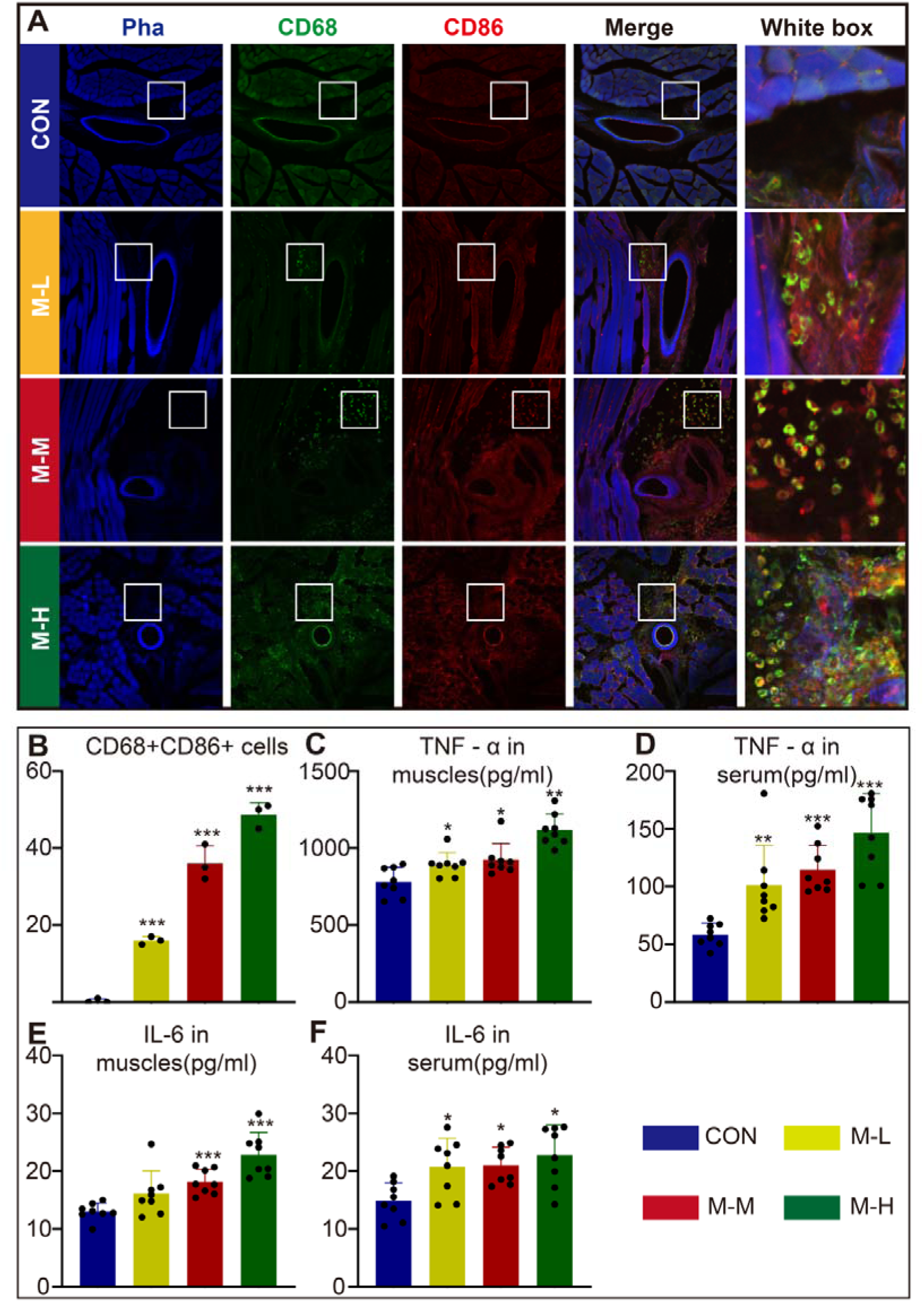
M1 macrophage polarization and pro-inflammatory cytokine levels in rats after eccentric contraction–induced muscle strain. (A) Representative immunofluorescence co-staining images of phalloidin (Pha, blue), CD68 (green), and CD86 (red) in the gastrocnemius muscle of each group; white boxes indicate the regions shown at higher magnification in the rightmost column. (B) Quantification of CD68□CD86□ double-positive (M1) cells in each group. (C, D) TNF-α content in the gastrocnemius muscle (C) and serum (D). (E, F) IL-6 content in the gastrocnemius muscle (E) and serum (F). n = 3 per group (A, B); data are presented as mean ± SD. n = 8 per group (C-F), data are presented as mean ± SEM.

## 4. Discussion

Utilizing a graded, parameter-controlled rat model of eccentric contraction-induced injury (comprising 50, 100, and 150 repetitions), this study elucidates that skeletal muscle functional impairment exacerbates in a manner dependent on the number of repetitions. This impairment is concomitant with microvascular rupture, erythrocyte extravasation, and disturbances in microcirculation. Extravasated erythrocytes may contribute to local iron accumulation through the heme–HO-1 pathway, accompanied by a ferroptosis-related molecular phenotype in skeletal muscle, as evidenced by the accumulation of Fe²□, depletion of GSH, increased levels of MDA, and diminished GPX4 content. Ferroptosis is accompanied by CD68□CD86□ M1 macrophage polarization and elevated levels of TNF-α and IL-6, these findings provide, to our knowledge, the first evidence supporting a model that connects microvascular rupture and erythrocyte extravasation with a potential ferroptosis–inflammation loop in eccentric skeletal muscle injury.

Conventional eccentric injury models, such as downhill running and electrical stimulation combined with pedal-driven stretch [10], offer limited control over the mechanical dose delivered to the muscle [11, 12]. In contrast, the Aurora servo-motor system provides precise regulation of stimulation parameters, joint angular velocity, and stretch amplitude, and has become a widely accepted standard for muscle mechanical assessment and modeling [13]. However, previous in vivo studies utilizing this system have predominantly focused on mouse hindlimb muscles or [14], in the case of rats, on submaximal training and rehabilitation protocols [15, 16]. The standardized rodent protocol developed by Lovering et al. involves a maximum of 30 stimulation cycles, which is insufficient to replicate the cumulative severe strain induced by high-frequency, long-duration exercise in clinical settings [13]. To address this limitation, the present study established a continuous stimulation–stretch gradient protocol comprising 50, 100, and 150 repetitions, with real-time recording of maximal eccentric torque. The torque exhibited the steepest decline within the first 100 repetitions, and the high-repetition group demonstrated significant lactate accumulation. The observed phenotypes-characterized by antalgic gait, asymmetrical weight-bearing, diminished endurance, and deep-tissue hyperalgesia-closely resembled clinical manifestations of muscle strain, thereby affirming the model’s face validity [15]. This robust and graded model serves as a valuable tool for analyzing dose-response relationships between the severity of injury and molecular events [7], and provides a platform for assessing therapeutic interventions [8].

A central mechanistic inquiry explored in this study pertains to the fate of extravasated erythrocytes and the consequent modifications in local heme metabolism. Eccentric exercise has been documented to disrupt capillary architecture, enhance vascular permeability, and impair microvascular reactivity [17]. In alignment with these findings, HE staining in the current investigation demonstrated microvascular disruption and substantial erythrocyte extravasation, while LSCI indicated significantly increased local perfusion. Severe eccentric injury, therefore, encompasses notable microvascular damage and microcirculatory disturbances beyond the destruction of myofibers [3, 17]. Extravasated erythrocytes undergo lysis within the interstitial space, releasing hemoglobin and heme, which likely constitute the primary sources of local heme accumulation [18]. Heme is predominantly cleared through hemopexin and macrophage-mediated heme degradation systems [19]. Although the cytotoxicity of heme is well supported in models of conditions such as intracerebral hemorrhage, hemolytic disorders, and atherosclerosis, it has been largely overlooked in the context of sports medicine [20]. HO-1, the rate-limiting enzyme in heme catabolism, is upregulated as a compensatory defensive mechanism. Nevertheless, the process functions as a double-edged sword, as heme degradation concurrently releases free iron, carbon monoxide, and biliverdin [21]. Under conditions of moderate stress, the upregulation of HO-1 serves a protective role [22]. Conversely, in scenarios where heme accumulation is extensive and persistent, it is possible that the iron derived from HO-1 may exceed the local capacities for storage, utilization, and export, thereby disrupting iron homeostasis [21, 22]. The application of multiplex immunofluorescence analysis introduces a spatial dimension to this understanding. HO-1-positive cells were found to cluster in perivascular regions and co-localize with CD68□ macrophages, while expression was sparse in parenchymal areas distant from vessels [21]. Previous studies suggest that macrophages are responsible for erythrophagocytosis and heme clearance, thus serving as the initial bearers of the local iron burden [18]. This cellular division of labor may elucidate why iron dyshomeostasis is characterized by an increased local tissue burden rather than a systemic overload, with muscle Fe²□ concentrations reaching approximately ten times those found in serum [18, 22]. Previous studies suggest that macrophages have the ability to export iron via ferroportin, and injured myofibers may enhance iron uptake through upregulation of transferrin receptor 1 (TfR1) [22]. One plausible explanation is that erythrocyte-derived heme undergoes catabolism by perivascular macrophages via HO-1, with the resultant iron being subsequently exported to adjacent myofibers. This iron transfer may create conditions conducive to iron-dependent lipid peroxidation and ferroptosis, as suggested in studies [20].

A pertinent mechanistic inquiry pertains to whether iron overload culminates in ferroptosis and the manner in which ferroptosis interacts with inflammation. In injured muscle, four indicators exhibited coordinated changes: Fe²□ accumulation, GSH depletion, elevated MDA levels, and suppressed GPX4 content. Collectively, these coordinated alterations represent a consistent molecular signature of ferroptosis [20]. Notably, these changes were confined to muscle tissue, as serum GSH levels and GPX4 content remained unaltered, highlighting the tissue-specific nature of this stress. Ferroptosis has been implicated in various conditions [23], including myocardial ischemia-reperfusion injury, acute kidney injury, and neurodegenerative diseases [24], and has also been reported in skeletal muscle contusion and musculoskeletal disorders [25, 26]. Nevertheless, its role in eccentric exercise–induced injury, characterized by microvascular rupture and redox imbalance, has not been thoroughly explored [18]. The current study expands the pathological context of ferroptosis to encompass this area [27]. Moreover, ferroptotic cells release DAMPs, with High Mobility Group Box 1 (HMGB1) being the most well-substantiated among them [28, 29]. Previous studies suggest that extracellular HMGB1 stimulates macrophage inflammatory responses through the AGER/RAGE pathway, facilitating M1-like polarization [30]. In contrast, previous studies suggest that TNF-α and IL-6 produced by M1 macrophages exert effects on adjacent myofibers [31, 32], and that TNF-α directly damages myoblasts and inhibits their differentiation [33]. Previous studies further suggest that pro-inflammatory signaling pathways lead to the upregulation of TfR1 via NF-κB-related mechanisms, resulting in increased cellular iron accumulation and potential impairment of iron export [34], which may enhance lipid peroxidation and susceptibility to ferroptosis [35]. Ferroptosis, in turn, is proposed to promote the release of DAMPs, activation of M1 macrophages, and secretion of cytokines, while these cytokines further disrupt myocellular iron homeostasis through autocrine and paracrine interactions [36]. One plausible explanation is that this mutual reinforcement creates a positive feedback loop, potentially elucidating the repetition-dependent worsening of functional impairment. These observations also align with the classical model in which the sequential polarization from M1 to M2 macrophages facilitates tissue repair. The infiltration of M1 macrophages during the acute phase is essential for the removal of necrotic debris [37]. However, previous studies suggest that a prolonged M1 response or a disrupted transition from M1 to M2 macrophages consistently hinders the regenerative process [38]. It is hypothesized that sustained myofiber ferroptosis perpetuates a pro-inflammatory microenvironment [39], thereby delaying the resolution of M1 activation and inhibiting the transition to pro-reparative phenotypes [40]. This proposed ferroptosis-induced persistence of M1 inflammation, coupled with an obstructed M1/M2 transition, may constitute a significant advancement in the understanding of muscle repair mechanisms, and may elucidate how a regulated cell death pathway could lead to regenerative failure through the dysregulation of immune response timing [41].

Several aspects of this study warrant further refinement. A ferroptosis-related phenotype was inferred from the coordinated changes in four biochemical markers: Fe²□ accumulation, GSH depletion, elevated MDA levels, and reduced GPX4 content [42], which align with its canonical features. MDA was measured using a TBA-based assay, which has limited analytical specificity. Future studies should incorporate more specific methods, such as LC–MS/MS quantification of F₂-isoprostanes, to further validate lipid peroxidation. Given that mitochondrial ultrastructural traits are not absolute prerequisites for ferroptosis, as different induction paradigms exhibit varying degrees of mitochondrial dependence [43], mitochondrial morphology was not utilized as a mandatory criterion in this study. Future research incorporating transmission electron microscopy and rescue experiments using ferroptosis-specific inhibitors (e.g., ferrostatin-1 and liproxstatin-1) would further substantiate the predominant role of ferroptosis and exclude other oxidative stress-related cell death modalities [44]. Additionally, it remains to be determined whether ferroptosis-driven sustained M1 polarization delays the transition to M2 and consequently impairs muscle regeneration [36–39]. This remains to be tested in future studies. Within this framework, the present study provides evidence that eccentric contraction-induced injury is associated with a microvascular rupture–erythrocyte extravasation-heme/iron overload pathway and a ferroptosis-related phenotype in skeletal muscle. This phenotype is accompanied by M1 macrophage polarization and elevated TNF-α and IL-6 levels, supporting a potential inflammatory feedback mechanism. Interventions targeting these mechanisms-such as iron chelation, ferroptosis inhibition, modulation of HO-1, or heme scavenging-could potentially offer novel, mechanism-based therapeutic strategies for addressing eccentric muscle injury.

## 5. Conclusions

This study demonstrated that a parameter-controlled eccentric contraction protocol induced graded skeletal muscle injury in rats, characterized by torque loss, gait abnormalities, hyperalgesia, and reduced endurance. These functional deficits were associated with microvascular disruption and erythrocyte extravasation, accompanied by heme accumulation, perivascular upregulation of HO-1, muscle iron accumulation, and a ferroptosis-related molecular phenotype. In parallel, M1 macrophage polarization and increased levels of TNF-α and IL-6 indicated activation of a pro-inflammatory response. Collectively, these findings support a potential heme–HO-1–iron–ferroptosis-associated inflammatory mechanism underlying eccentric muscle injury. Future translational studies targeting heme scavenging, ferroptosis inhibition, or inflammatory modulation may help establish causal relationships and facilitate the development of mechanism-based therapeutic strategies for eccentric skeletal muscle injury.

## Author Contributions

Conceptualization, ZW.L., JS.Y. and YY.W.; methodology, XY.W., JS.Y. and YY.W.; software, MH.L.; validation, MH.L.; formal analysis, MH.L.; investigation, XY.W., ZW.L., FM.Y., Q.S., M.Z., S.C. and YT.J.; resources, XY.W.; data curation, MH.L., FM.Y., Q.S., M.Z., S.C. and YT.J.; writing—original draft preparation, MH.L.; writing—review and editing, XY.W.; visualization, MH.L.; supervision, XY.W.; project administration, JS.Y. and YY.W.; funding acquisition, JS.Y. and YY.W. All authors have read and agreed to the published version of the manuscript.

## Funding

This work was supported by the National Natural Science Foundation of China (82675980) and the China Academy of Chinese Medical Sciences Innovation Fund (ID: CI2025C001).

## Institutional Review Board Statement

The animal study protocol was approved by the Ethics Committee of the Institute of Acupuncture and Moxibustion, China Academy of Chinese Medical Sciences (Beijing, China) (approval number: D2024-01-26-05). All procedures were conducted in accordance with the relevant institutional and national guidelines for the care and use of laboratory animals, the ARRIVE guidelines, and the 3Rs principles (replacement, reduction, and refinement). Male Wistar rats weighing 180 to 220 g were purchased from the Institute of Laboratory Animal Science, Chinese Academy of Medical Sciences and were maintained under a standard 12-hour light–dark cycle with free access to food and water. The study was not designed to evaluate sex-specific differences in eccentric contraction-induced muscle injury; therefore, only male rats were used.

## Informed Consent Statement

Not applicable.

## Data Availability Statement

The data presented in this study are available on request from the corresponding author.

## Acknowledgments

Not applicable.

## Conflicts of Interest

The authors declare no conflicts of interest.

## Notes

### Competing Interest Statement

The authors have declared no competing interest.

